## Supplementary Tables and Figures for "Structural basis for recognition of two HLA-A2-restricted SARS-CoV-2 spike epitopes by public and private T cell receptors"

Daichao Wu et al.

**Supplementary Tables 1–11**

**Supplementary Figures 1–4**

### Supplementary Table 1. SARS-CoV2-specific TCR germline genes and CDR3 sequences

| Receptor | α chain | | | | β chain | | | |
|---|---|---|---|---|---|---|---|---|
| | **TRAV** | **CDR3α** | **TRAJ** | **Patients from (26)** | **TRBV** | **CDR3β** | **TRBJ** | **Patients from (26)** |
| **RLQ3** | TRAV16 | CALSGFNNAGNMLTF | TRAJ39 | p1434 | TRBV11-2 | CASSLGGAGGADTQYF | TRBJ2-3 | p1434 |
| **RLQ5** | TRAV12-2 | CALSSNDYKLSF | TRAJ20 | p1437<br>p1499 | TRBV6-5 | CATTERDQETQYF | TRBJ2-5 | p1437 |
| **RLQ7** | TRAV38-2DV8 | CASSGNTPLVF | TRAJ29 | p1445 | TRBV12-3 | CASTWGRASTDTQYF | TRBJ2-3 | p1445 |
| **RLQ8** | TRDV1 | CALGERDSWGKFQF | TRAJ24 | p1448 | TRBV20-1 | CSALTPALAGGVPETQYF | TRBJ2-5 | p1448 |
| **YLQ7** | **TRAV12-2** | CAVNRDDKIIF | TRAJ30 | p1434<br>p1437<br>p1445<br>p1449<br>p1480<br>p1484<br>p1495 | TRBV7-9 | CASSPDIEQYF | TRBJ2-7 | p1434<br>p1437<br>p1445<br>p1480<br>p1484 |

**Supplementary Table 2. Affinity of TCRs for SARS-CoV-2 spike epitopes and epitope variants**

| **Virus** | **Epitope** | **Identity with RLQ/YLQ** | **Sequence** | **YLQ7** | **RLQ3** | **RLQ5** | **RLQ7** | **RLQ8** |
|---|---|---|---|---|---|---|---|---|
| SARS-CoV-2 | RLQ | 9/9 | RLQSLQTYV | | 32.9 μM | 3.4 μM | 66.4 μM | 9.7 μM |
| MERS | RLT | 4/9 | RLTTLNAFV | | NB | | | |
| SARS | RLQ | 9/9 | RLQSLQTYV | | 32.9 μM | | | |
| HKU1 | RLT | 5/9 | RLTALNAYV | | NB | | | |
| OC43 | RLT | 5/9 | RLTALNAYV | | NB | | | |
| NL63 | RLA | 4/9 | RLAALNAFV | | NB | | | |
| 229E | RLA | 4/9 | RLAALNVFV | | NB | | | |
| SARS-CoV-2 variant | T1006I | 8/9 | RLQSLQ**I**YV | | 121 μM | 23.0 μM | 92.9 μM | 122 μM |
| SARS-CoV-2 | YLQ | 9/9 | YLQPRTFLL | 1.8 μM | | | | |
| MERS | KLQ | 7/9 | KLQPLTFLL | NB | | | | |
| SARS | YLK | 6/9 | YLKPTTFML | NB | | | | |
| HKU1 | PLS | 4/9 | PLSKRQYLL | NB | | | | |
| OC43 | PLT | 4/9 | PLTSRQYLL | NB | | | | |
| SARS-CoV-2 variant | P272L | 8/9 | YLQ**L**RTFLL | 127 μM | | | | |

Affinities were measured by SPR. NB, no binding.

**Supplementary Table 3. YLQ and RLQ epitope variants in SARS-CoV-2 sequences**

| Substitution | Sequence | Frequency[1] |
|---|---|---|
| P272L | YLQ**L**RTFLL | 0.59% |
| T1006I | RLQSLQ**I**YV | 0.04% |
| R273S | YLQP**S**TFLL | 0.011% |
| T274I | YLQPR**I**FLL | 0.007% |
| R273K | YLQP**K**TFLL | 0.006% |
| P272H | YLQ**H**RTFLL | 0.006% |
| P272S | YLQ**S**RTFLL | 0.005% |
| T274N | YLQPR**N**FLL | 0.003% |
| Q271K | YL**K**PRTFLL | 0.003% |
| Q1005H | RLQSL**H**TYV | 0.003% |
| L270F | Y**F**QPRTFLL | 0.0002% |

[1]Frequency of substitution in SARS-CoV-2 spike glycoprotein sequences in the GISAID database (www.gisaid.org).

### Supplementary Table 4. Data collection and refinement statistics

| | RLQ3 | RLQ–HLA-A2 | RLQ3–RLQ–HLA-A2 | YLQ7 | YLQ–HLA-A2 | YLQ7–YLQ–HLA-A2 |
|---|---|---|---|---|---|---|
| PDB | 7N1C | 7N1B | 7N1E | 7N1D | 7N1A | 7N1F |
| **Data collection** | | | | | | |
| Resolution range (Å) | 46.6–1.88 (1.95–1.88) | 43.2–2.81 (2.91–2.81) | 49.4–2.30 (2.38–2.30) | 38.8–2.35 (2.43–2.35) | 43.3–2.06 (2.14–2.06) | 48.6–2.39 (2.48–2.39) |
| Space group | *P*212121 | *P*1 | *P*1211 | *I*4 | *P*3212 | *C*121 |
| Unit cell parameters | 42.1, 101.9, 104.7 90, 90, 90 | 47.2, 49.3, 117.1 91.9, 92.4, 118.6 | 52.8, 68.2, 148.6 90, 94.6, 90 | 109.8, 109.8, 99.6 90, 90, 90 | 85.1, 85.1, 214.2 90, 90, 120 | 225.3, 49.1, 91.5 90, 91.8, 90 |
| Total reflections[a] | 239,229 (21,569) | 37,697 (3,276) | 159,246 (15,350) | 172,549 (17,833) | 558,646 (56,058) | 133,860 (12,215) |
| Unique reflections[a] | 37,372 (3,632) | 21,652 (2,067) | 46,769 (4,435) | 24,641 (2,476) | 54,889 (5,464) | 39,704 (3,697) |
| Multiplicity[a] | 6.4 (5.9) | 1.7 (1.6) | 3.4 (3.5) | 7.0 (7.2) | 10.2 (10.3) | 3.4 (3.3) |
| Completeness (%)[a] | 99.8 (98.6) | 96.1 (89.1) | 99.2 (94.8) | 99.9 (100.0) | 100 (99.9) | 99.0 (92.4) |
| Mean $I/\sigma(I)$[a] | 18.6 (2.9) | 9.4 (5.8) | 12.6 (3.0) | 12.8 (2.9) | 17.8 (5.6) | 9.4 (2.1) |
| Wilson *B* factor (Å$^2$) | 32.6 | 25.6 | 32.2 | 44.9 | 28.6 | 48.3 |
| $R_{merge}$[a,b] | 0.067 (0.399) | 0.128 (0.320) | 0.148 (0.472) | 0.108 (0.549) | 0.141 (0.427) | 0.065 (0.363) |
| CC1/2 | 0.996 (0.937) | 0.911 (0.638) | 0.952 (0.753) | 0.994 (0.832) | 0.991 (0.958) | 0.993 (0.938) |
| **Refinement** | | | | | | |
| Reflections used in refinement[a] | 37,368 (3,631) | 21,643 (2,066) | 46,625 (4,411) | 24,640 (2,476) | 54,879 (5,464) | 39,630 (3,680) |
| $R_{work}$[c] | 0.201 (0.256) | 0.201 (0.242) | 0.213 (0.277) | 0.204 (0.265) | 0.196 (0.226) | 0.199 (0.285) |
| $R_{free}$[c] | 0.231 (0.280) | 0.271 (0.340) | 0.259 (0.369) | 0.271 (0.377) | 0.236 (0.280) | 0.235 (0.362) |
| No. of protein atoms | 3,462 | 6,261 | 6,631 | 3,473 | 6,289 | 6,567 |
| No. of waters | 267 | | 253 | 142 | 482 | 87 |
| Protein residues | 440 | 764 | 826 | 439 | 766 | 818 |
| r.m.s.d. from ideality | | | | | | |
| Bond lengths (Å) | 0.009 | 0.010 | 0.010 | 0.008 | 0.009 | 0.009 |
| Bond angles (º) | 1.44 | 1.40 | 1.30 | 1.30 | 1.25 | 1.26 |
| Ramachandran plot statistics | | | | | | |
| Favored (%) | 96.1 | 97.1 | 95.1 | 95.9 | 97.6 | 96.2 |
| Allowed (%) | 3.7 | 2.8 | 4.7 | 3.9 | 2.4 | 3.5 |
| Disallowed (%) | 0.2 | 0.1 | 0.3 | 0.2 | 0.0 | 0.4 |
| Rotamer outliers (%) | 1.9 | 0.5 | 0.8 | 0.3 | 0.9 | 1.6 |
| Clashscore | 6.0 | 11.0 | 9.3 | 8.9 | 7.7 | 8.0 |
| Average *B* factor (Å$^2$) | 41.4 | 30.7 | 41.1 | 47.5 | 35.4 | 62.6 |
| Protein | 41.3 | 30.7 | 41.2 | 47.5 | 35.2 | 62.7 |
| Waters | 42.2 | | 40.5 | 47.6 | 38.0 | 53.4 |

[a]Values in parentheses correspond to the highest resolution shell.

[b]$R_{merge} = \sum|I_j - \langle I \rangle|/\sum I_j$, where $I_j$ is the intensity of an individual reflection and $\langle I \rangle$ is the average intensity of that reflection.

[c]$R_{work}$ ($R_{free}$) $= \sum||F_o| - |F_c||/\sum|F_o|$; 5.0% of data were used for $R_{free}$.

### Supplementary Table 5. Interactions between RLQ3 and HLA-A2

| HLA-A2 | RLQ3 | | |
|---|---|---|---|
| | Hydrogen bonds | Van der Waals contacts | Water bridges |
| α1 | | | |
| K66H | | N92α(2) | |
| V76H | | N49β(3), | |
| T80H | N49β(Nδ2) T80H(Oγ1), | N49β(2), | |
| α2 | | | |
| H145H | | L94β(2) | |
| K146H | | G26β(1), L94β(3), G96β(2) | G95β(N) S253 K146H(O), G96β(N) S253 K146H(O), D101β(Oδ2) S253 K146H(O) |
| W147H | G96β(O) W147H(Nε1), | G96β(3) | |
| A149H | | L94β(1), D101β(2) | |
| A150H | R48α(Nη2) A150H(O), | R48α(3), A100β(1), D101β(5) | A100β(O) S71 A150H(O), |
| H151H | S51α(Oγ) H151H(Nε2), | I50α(1), S51α(4) | |
| E154H | | I50α(1) | |
| Q155H | E31α(Oε2) Q155H(Nε2), R48α(Nη1) Q155H(Nε2), | E31α(3), R48α(3), I50α(3), F91α(4) | P30α(O) S68 Q155H(Oε1, Nε2), E31α(Oε1) S68 Q155H(Oε1, Nε2) |

Contact residues were identified with the CONTACT program (69). Hydrogen bonds were calculated using a cut-off distance of 3.5 Å. The cut-off distance for van der Waals contacts was 4 Å.

**Supplementary Table 6. Interactions between RLQ3 and RLQ peptide**

| RLQ | RLQ3 | | |
|---|---|---|---|
| | Hydrogen bonds | Van der Waals contacts | Water bridges |
| S4p | N92α(Nδ2) S4p(Oγ) | S29α(4), F91α(2) N92α(2) | |
| L5p | | F91α(9), N96α(2) A100β(1) | |
| Q6p | F91α(O) Q6p(Nε2), N93α(O) Q6p(Nε2), G95α(N) Q6p(Oε1), N96α(Nδ2) Q6p(N), N96α(Oδ1) Q6p(N), N96α(Nδ2) Q6p(O), N96α(Oδ1) Q6p(Nε2), | F91α(2), N93α(3), A94α(2), G95α(3), N96α(9) A97β(3) | |
| T7p | | G96β(4) | |
| Y8p | G96β(O) Y8p(N), G96β(O) Y8p(O), | Q48β(11), V53β(1), G95β(3), G96β(6), G98β(2), | Y8p(Oη) S106 Q46β(Nε2), Y8p(Oη) S106 Q48β(Oε1) |

Contact residues were identified with the CONTACT program (69). Hydrogen bonds were calculated using a cut-off distance of 3.5 Å. The cut-off distance for van der Waals contacts was 4 Å.

**Supplementary Table 7. Predicted TCR binding energy changes from computational alanine scanning of peptide positions in SARS-CoV-2 TCR–pMHC complexes.**

| **RLQ3–RLQ–HLA-A2** | | **YLQ7–YLQ–HLA-A2** | |
|---|---|---|---|
| **Mutation** | **ΔΔ*G*[1]** | **Mutation** | **ΔΔ*G*[1]** |
| R1A | 0 | Y1A | 0 |
| L2A | 0 | L2A | 0 |
| Q3A | 0 | **Q3A** | **1.3** |
| S4A | -0.1 | P4A | 0.1 |
| **L5A** | **1.0** | **R5A** | **3.0** |
| **Q6A** | **2.3** | **T6A** | **1.2** |
| T7A | 0.2 | F7A | 0.2 |
| **Y8A** | **1.9** | L8A | 0.7 |
| V9A | 0 | L9A | 0 |

[1]TCR binding energy changes for individual peptide residue alanine mutations, calculated using Rosetta with experimentally determined TCR–pMHC complex structures as input. Values are in Rosetta Energy Units (REU), which correspond to energy in kcal/mol. Substitutions with high predicted ΔΔ*G*s (≥ 1 REU), reflective of likely energetic hotspots, are in bold.

**Supplementary Table 8. Predicted HLA-A2 binding affinities and TCR binding effects for epitope variants across coronaviruses and within SARS-CoV-2**

| Lineage[1] | Accession ID[2] | Virus/ Variant | Identity[3] | Sequence | Pred. HLA-A2 affinity (nM)[4] | Predicted TCR ΔΔG[5] | Measured TCR ΔΔG, kcal/mol[6] |
|---|---|---|---|---|---|---|---|
| *RLQ epitope orthologs* | | | | | | | |
| B, clade 1b | QHD43416.1 | SARS-CoV-2 | 9 | RLQSLQTYV | 11.92 | 0 | |
| B, clade 1b | QHR63300.2 | RaTG13 | 9 | RLQSLQTYV | 11.92 | 0 | |
| B, clade 1b | QIG55945.1 | GD_pangolin | 9 | RLQSLQTYV | 11.92 | 0 | |
| B, clade 1a | AYV99761.1 | SARS-CoV | 9 | RLQSLQTYV | 11.92 | 0 | |
| B, clade 1a | QJE50589.1 | SHC014 | 9 | RLQSLQTYV | 11.92 | 0 | |
| B, clade 1a | AGZ48828.1 | WIV1 | 9 | RLQSLQTYV | 11.92 | 0 | |
| B, clade 1a | QND76034.1 | HKU3 | 9 | RLQSLQTYV | 11.92 | 0 | |
| B, clade 1a | AHX37569.1 | LYRa3 | 9 | RLQSLQTYV | 11.92 | 0 | |
| B, clade 2 | EPI_ISL_412977 | RmYN02 | 9 | RLQSLQTYV | 11.92 | 0 | |
| B, clade 2 | ATO98120.1 | rs4081 | 9 | RLQSLQTYV | 11.92 | 0 | |
| B, clade 2 | ABD75323.1 | Rf1 | 9 | RLQSLQTYV | 11.92 | 0 | |
| B, clade 3 | APO40579.1 | BtKY72 | 9 | RLQSLQTYV | 11.92 | 0 | |
| B, clade 3 | YP_003858584.1 | BM48-31 | 9 | RLQSLQTYV | 11.92 | 0 | |
| A | YP_173238.1 | HKU1 | 5 | RLTALNAYV | 8.27 | 1.6 | NB |
| A | ANZ78843.1 | OC43 | 5 | RLTALNAYV | 8.27 | 1.6 | |
| A | ACN89742.1 | MHV-1 | 5 | RLTALNAYV | 8.27 | 1.6 | |
| A | YP_009113025.1 | HKU24 | 5 | RLTALNAYV | 8.27 | 1.6 | |
| C | YP_009047204.1 | MERS | 4 | RLTTLNAFV | 13.32 | 1.6 | |
| C | YP_001039953.1 | HKU4 | 5 | RLISLNAFV | 4.97 | 2.2 | NB |
| C | YP_001039962.1 | HKU5 | 5 | RLTSLNAFV | 10.18 | 2.2 | |
| C | AGC51116.1 | KW2E-F93 | 4 | RLTTLNAFV | 13.32 | 1.6 | |
| C | QGA70692.1 | HKU31 | 5 | RLTSLNAFV | 10.18 | 2.2 | |
| D | YP_001039971.1 | HKU9 | 5 | RMMVLNTYV | 3.76 | 1.5 | |
| D | ADX59466.1 | KY24 | 5 | RMVVLNTYV | 21.45 | 1.6 | |
| D | QDF43840.1 | GX2018 | 5 | RTVVLNTYV | 841.78 | 1.6 | |
| Hibecovirus | YP_009072440.1 | Zhejiang2013 | 7 | RLQVLQTFV | 36.54 | 0.8 | NB |
| alpha | AFV53148.1 | NL63 | 4 | RLAALNAFV | 5.36 | 2.2 | NB |
| alpha | ABB90529.1 | 229E | 4 | RLAALNVFV | 5.01 | 2.1 | |
| *YLQ epitope orthologs* | | | | | | | |
| B, clade 1b | QHD43416.1 | SARS-CoV-2 | 9 | YLQPRTFLL | 4.3 | 0 | |
| B, clade 1b | QHR63300.2 | RaTG13 | 9 | YLQPRTFLL | 4.3 | 0 | |
| B, clade 1b | QIG55945.1 | GD_pangolin | 7 | YLAPRTFML | 3.01 | 1.9 | 1.7 |
| B, clade 1a | AYV99761.1 | SARS-CoV | 6 | YLKPTTFML | 12.25 | 4.6 | NB |
| B, clade 1a | QJE50589.1 | SHC014 | 6 | YLKPTTFML | 12.25 | 4.6 | NB |
| B, clade 1a | AGZ48828.1 | WIV1 | 6 | YLKPTTFML | 12.25 | 4.6 | |

| | | | | | | | |
|---|---|---|---|---|---|---|---|
| B, clade 1a | QND76034.1 | HKU3 | 4 | NLKYSTFML | 616.38 | 3.5 | |
| B, clade 1a | AHX37569.1 | LYRa3 | 6 | YLKPTTFML | 12.25 | 4.6 | NB |
| B, clade 2 | ATO98120.1 | rs4081 | 4 | NLKYSTFML | 616.38 | 3.5 | |
| B, clade 2 | ABD75323.1 | Rf1 | 4 | NLKQSTFML | 756.46 | 3.2 | |
| B, clade 2 | EPI_ISL_412977 | RmYN02 | 3 | SLKLTTIML | 1121.94 | 4.2 | |
| B, clade 3 | APO40579.1 | BtKY72 | 4 | HLKPLTMLA | 1682.36 | 3.5 | |
| B, clade 3 | YP_003858584.1 | BM48-31 | 4 | HLKPLTMLV | 157.7 | 3.5 | |
| A | YP_173238.1 | HKU1 | 4 | PLSKRQYLL | 7176.36 | 1 | |
| A | ANZ78843.1 | OC43 | 4 | PLTSRQYLL | 5129.13 | 1.8 | |
| A | ACN89742.1 | MHV-1 | 3 | PLVERQYLF | 12445.47 | 0.6 | |
| A | YP_009113025.1 | HKU24 | 4 | PLIKREYLL | 2115.53 | 1.6 | |
| C | YP_009047204.1 | MERS | 7 | KLQPLTFLL | 6.12 | 2.3 | NB |
| C | YP_001039953.1 | HKU4 | 4 | KLHQLTYLL | 14.83 | 4.1 | |
| C | YP_001039962.1 | HKU5 | 5 | KLHPLTYLL | 9.27 | 5.1 | |
| C | AGC51116.1 | KW2E-F93 | 4 | YLYELPYLL | 1.93 | 12.8 | |
| C | QGA70692.1 | HKU31 | 2 | QLHKLNYLV | 59.62 | 5.2 | |
| D | YP_001039971.1 | HKU9 | 3 | HLINRDLLV | 49.64 | 2.3 | |
| D | ADX59466.1 | KY24 | 2 | PLSKQDVLV | 6784.7 | 4.2 | |
| D | QDF43840.1 | GX2018 | 2 | PIVQRELLV | 17412.75 | 1.5 | |
| Hibecovirus | YP_009072440.1 | Zhejiang2013 | 3 | QLKKSTFMF | 14082.76 | 3.1 | |
| alpha | AFV53148.1 | NL63 | 1 | AFATFVDVL | 10799.6 | 5.8 | NB |
| alpha | ABB90529.1 | 229E | 2 | ALASYADVL | 147.99 | 9 | NB |
| *SARS-CoV-2 Variants* | | | | | | | |
| - | - | T1006I | 8 | RLQSLQIYV | 6.57 | 0.1 | 0.77 |
| - | - | Q1005H | 8 | RLQSLHTYV | 14.92 | 1.3 | |
| - | - | L270F | 8 | YFQPRTFLL | 1114.21 | 0 | |
| - | - | P272L | 8 | YLQLRTFLL | 9.56 | -0.2 | 2.5 |
| - | - | T274N | 8 | YLQPRNFLL | 4.73 | 1.2 | |
| - | - | T274I | 8 | YLQPRIFLL | 3.21 | 1.3 | |
| - | - | R273S | 8 | YLQPSTFLL | 3.56 | 2.6 | |
| - | - | Q271K | 8 | YLKPRTFLL | 16.03 | 1.0 | |
| - | - | R273K | 8 | YLQPKTFLL | 4 | 2.9 | |
| - | - | P272H | 8 | YLQHRTFLL | 5.65 | 0.0 | |
| - | - | P272S | 8 | YLQSRTFLL | 4.65 | -0.2 | |

[1]Betacoronavirus lineage (A-D, or Hibecovirus), or "alpha" in the case of alphacoronaviruses. For lineage B (sarbecoviruses), which includes SARS-CoV-2 and SARS-CoV, clade information is also provided. "-": SARS-CoV-2 variant.
[2]NCBI Genbank or GISAID accession ID.
[3]Number of residues identical with SARS-CoV-2 YLQ or RLQ epitope.
[4]Predicted HLA-A*02:01 binding ($IC_{50}$, nM), from NetMHCPan 4.1 (50).
[5]Predicted RLQ3 or YLQ7 TCR binding affinity change ($\Delta\Delta G$) for altered peptide versus RLQ or YLQ epitope, calculated using Rosetta (48) with RLQ3-RLQ-HLA-A2 or YLQ7-YLQ-HLA-A2 complex structures. Units are Rosetta Energy Units, which are comparable to kcal/mol.

[6]Measured RLQ3 or YLQ7 TCR binding affinity change versus RLQ or YLQ epitope, from this study. NB: no measurable binding detected.

**Supplementary Table 9. Interactions between YLQ7 and HLA-A2**

| HLA-A2 | YLQ7 | | |
|---|---|---|---|
| | Hydrogen bonds | Van der Waals contacts | Water bridges |
| α1 | | | |
| R65H | R65H(Nε) D94α(Oδ2) | D94α(4) | |
| K66H | | D94α(1) | |
| A69H | | | A69H(O) S66 Q50β(Nε2) |
| Q72H | | L55β(3) | |
| T73H | | R31β(1), Q50β(2) | T73H(Oγ1) S66 Q50β(Nε2) |
| V76H | | N51β(1) | |
| α2 | | | |
| A150H | | | A150H(O) S53 Y51α(Oη), |
| H151H | | F49α(4), | |
| E154H | S52α(Oγ) E154H(Oε2) | Y51α(3), S52α(4) | E154H (Oε2) S41 Y51α(N)<br>E154H (O) S16 S52α(Oγ) |
| Q155H | Q31α(Oε1) Q155H(O),<br>S32α(Oγ) Q155H(Oε1), | Q31α(1), S32α(2), Y51α(10) I98β(1) | |
| R157H | S52α(Oγ) R157H(Nη2) | S52α(2) | R157H (NH2) S16 S52α(Oγ) |
| A158H | | Y51α(1), K67α(1) | |
| Y159H | | Q31α(1) | |
| E166H | R28α(Nε) E166H(Oε1),<br>R28α(Nε) E166H(Oε2), | R28α(5) | |

Contact residues were identified with the CONTACT program (69). Hydrogen bonds were calculated using a cut-off distance of 3.5 Å. The cut-off distance for van der Waals contacts was 4 Å.

**Supplementary Table 10. Interactions between YLQ7 and YLQ peptide**

| YLQ | YLQ7 | | |
|---|---|---|---|
| | Hydrogen bonds | Van der Waals contacts | Water bridges |
| Y1p | | D27α(1), G29α(1) | |
| Q3p | Q31α(Oε1) Qp3(Nε2) | Q31α(4) | |
| P4p | | G29α(1), Q31α(2) D94α(5) | P4p(O) S35 D95α(Oδ2) |
| R5p | Q31α(Nε2) R5p(N) S32α(Oγ) R5p(Nε) D95α(Oδ1) R5p(Nη1) D97β(Oδ1) R5p(Nη2) D97β(Oδ2) R5p(Nη2) D97β(Oδ1) R5p(Nη1) | Q31α(2) S32α(1), N92α(2), D94α(2) D95α(5) R31β(1) D97β(6) I98β(8) | |
| T6p | D95α(Oδ1) T6p(N), D95α(Oδ2) T6p(N), D95α(Oδ1) T6p(O), D95α(Oδ2) T6p(Oγ1) R31β(Nη1) T6p(O), R31β(Nη1) T6p(Oγ1), R31β(Nη2) T6p(O), D97β(Oδ1) T6p(O) | D95α(9) R31β(2) | |
| F7p | | D97β(4) | |
| L8p | | N30β(1) R31β(2) Q50β(3) | |

Contact residues were identified with the CONTACT program (69). Hydrogen bonds were calculated using a cut-off distance of 3.5 Å. The cut-off distance for van der Waals contacts was 4 Å.

**Supplementary Table 11. Sequences of TCR genes codon-optimized for expression in *E. coli***

| TCR gene name | Nucleotide sequence |
|---|---|
| RLQ3<br>alpha | CAT**ATG**CAGCGTGTGACCCAGCCGGAAAAACTGCTGAGCGTGTTTAAAGGCGCGCCGGTTGAACT<br>GAAATGCAATTATAGCTATAGCGGCAGCCCGGAACTGTTTTGGTATGTGCAGTATAGCCGCCAGC<br>GCCTGCAGCTGTTGCTGCGCCATATTAGCCGTGAAAGCATTAAAGGCTTTACCGCGGATCTGAAC<br>AAAGGCGAAACCAGCTTTCATCTGAAAAAACCGTTTGCGCAGGAAGAGGATAGCGCGATGTATTA<br>CTGCGCGCTGAGCGGCTTTAACAATGCGGGTAACATGCTGACCTTTGGTGGCGGTACCCGTCTGA<br>TGGTGAAACCGAACATTCAGAACCCGGATCCGGCGGTTTATCAGCTGCGTGATAGCAAAAGCAGC<br>GATAAAAGCGTGTGCCTGTTTACCGATTTTGATAGCCAGACCAACGTGAGCCAGAGCAAAGATAG<br>CGATGTGTATATTACCGATAAATGCGTGCTGGATATGCGCAGCATGGATTTTAAAAGCAACAGCG<br>CGGTGGCGTGGAGCAACAAAAGCGATTTTGCGTGCGCGAACGCGTTTAACAACAGCATCATTCCG<br>GAAGATACCTTTTTCCCGAGCCCGGAAAGCAGC**TAATGA**CTCGAG |
| RLQ3<br>beta | CAT**ATG**GGTGTTGCGCAGAGCCCGCGCTATAAAATCATTGAAAAACGTCAGAGCGTGGCCTTTTG<br>GTGCAACCCGATTAGCGGCCACGCGACCCTGTATTGGTATCAACAGATTCTGGGCCAGGGTCCGA<br>AACTGCTGATTCAGTTTCAGAACAATGGCGTGGTTGATGACAGCCAGCTGCCGAAAGATCGCTTT<br>AGCGCGGAACGTCTGAAAGGCGTGGATAGCACCCTGAAAATCCAGCCGGCCAAACTGGAAGATAG<br>CGCGGTGTATCTGTGTGCGAGCAGCCTGGGCGGTGCGGGTGGCGCCGATACCCAGTACTTTGGTC<br>CGGGTACCCGTCTGACCGTGCTGGAAGATCTGAAAAACGTGTTTCCGCCGGAAGTGGCGGTGTTT<br>GAACCGAGCGAAGCGGAAATTAGCCATACCCAGAAAGCGACCCTGGTGTGCCTGGCGACCGGTTT<br>TTATCCGGATCATGTGGAACTGAGCTGGTGGGTGAACGGCAAAGAAGTGCATAGCGGCGTGTGCA<br>CCGATCCGCAGCCGCTGAAAGAACAGCCGGCGCTGAACGATAGCCGTTATGCGCTGAGCAGCCGT<br>CTGCGTGTGAGCGCGACCTTTTGGCAGAACCCGCGCAACCATTTTCGCTGCCAAGTTCAGTTTTA<br>CGGACTGTCGGAAAACGATGAATGGACCCAGGATCGTGCGAAACCGGTGACCCAGATTGTGAGCG<br>CGGAAGCGTGGGGCCGCGCGGAT**TAATGA**CTCGAG |
| YLQ7<br>alpha | CAT**ATG**CAGAAAGAGGTGGAACAAAACAGCGGTCCGCTGAGCGTTCCGGAGGGTGCGATTGCGAG<br>CCTGAACTGCACCTACAGCGATCGTGGTAGCCAGAGCTTCTTTTGGTACCGTCAATATAGCGGCA<br>AAAGCCCGGAGCTGATCATGTTCATTTATAGCAACGGTGACAAGGAAGATGGCCGTTTTACCGCG<br>CAGCTGAACAAAGCGAGCCAATACGTGAGCCTGCTGATTCGTGACAGCCAGCCGAGCGATAGCGC<br>GACCTATCTGTGCGCGGTTAACCGTGACGATAAGATCATTTTCGGTAAAGGCACCCGTCTGCACA<br>TCCTGCCGAACATTCAGAACCCGGACCCGGCGGTGTACCAACTGCGTGACAGCAAGAGCAGCGAT<br>AAAAGCGTGTGCCTGTTCACCGACTTTGATAGCCAGACCAACGTTAGCCAAAGCAAGGACAGCGA<br>TGTGTATATCACCGACAAATGCGTTCTGGATATGCGTAGCATGGACTTTAAGAGCAACAGCGCGG<br>TTGCGTGGAGCAACAAAAGCGATTTCGCGTGCGCGAACGCGTTTAACAACAGCATCATTCCGGAG<br>GACACCTTCTTTCCGAGCCCGGAAAGCAGC**TAATGA**CTCGAG |
| YLQ7<br>beta | CAT**ATG**GACACCGGTGTGAGCCAGAACCCGCGTCACAAGATCACCAAACGTGGCCAAAACGTTAC<br>CTTCCGTTGCGATCCGATTAGCGAGCACAACCGTCTGTACTGGTATCGTCAGACCCTGGGTCAAG<br>GCCCGGAATTCCTGACCTACTTTCAGAACGAGGCGCAACTGGAAAAGAGCCGTCTGCTGAGCGAC<br>CGTTTCAGCGCGGAGCGTCCGAAAGGTAGCTTTAGCACCCTGGAGATCCAGCGTACCGAACAAGG<br>CGACAGCGCGATGTACCTGTGCGCGAGCAGCCCGGATATTGAGCAGTATTTTGGTCCGGGTACCC<br>GTCTGACCGTGACCGAAGACCTGAAGAACGTTTTCCCGCCGGAAGTGGCGGTTTTTGAACCGAGC<br>GAGGCGGAAATCAGCCACACCCAAAAAGCGACCCTGGTGTGCCTGGCGACCGGTTTTTATCCGGA<br>TCACGTGGAGCTGTCCTGGTGGGTTAACGGCAAGGAAGTGCATAGCGGCGTTTGCACCGACCCGC<br>AGCCGCTGAAAGAGCAACCGGCGCTGAACGATAGCCGTTATGCGCTGAGCAGCCGTCTGCGTGTG<br>AGCGCGACCTTTTGGCAGAACCCGCGTAACCACTTCCGTTGCCAGGTTCAATTTTATGGTCTGAG<br>CGAGAACGACGAATGGACCCAGGATCGTGCGAAGCCGGTGACCCAAATTGTTAGCGCGGAAGCGT<br>GGGGCCGTGCGGAT**TAATGA**CTCGAG |

Start and stop codons are shown in bold. Sequences coding for upstream (Nde I) and downstream (Xho I) cloning sites are underlined.

### Supplementary Figure Legends

**Figure S1.** SPR analysis of RLQ-specific TCRs binding to RLQ epitopes and T1006I variant. **(a)** (upper) TCR RLQ5 at concentrations of 0.39, 0.78, 1.56, 3.12, 6.25, 12.5, 25, and 50.0 μM was injected over immobilized RLQ–HLA-A2 (1200 RU). (lower) Fitting curve for equilibrium binding that resulted in a $K_D$ of 3.4 μM. **(b)** (upper) TCR RLQ5 at concentrations of 0.39, 0.78, 1.56, 3.12, 6.25, 12.5, 25, and 50.0 μM was injected over immobilized T1006I–HLA-A2 (1200 RU). (lower) Fitting curve for equilibrium binding that resulted in a $K_D$ of 23.0 μM. **(c)** (upper) TCR RLQ7 at concentrations of 1.25, 2.5, 5, 10, 25, 20.0, and 40 μM was injected over immobilized RLQ–HLA-A2 (1200 RU). (lower) Fitting curve for equilibrium binding that resulted in a $K_D$ of 66.4 μM. **(d)** (upper) TCR RLQ7 at concentrations of 0.39, 0.78, 1.56, 3.12, 6.25, 12.5, 25, and 50.0 μM was injected over immobilized T1006I–HLA-A2 (1200 RU). (lower) Fitting curve for equilibrium binding that resulted in a $K_D$ of 92.9 μM. **(e)** (upper) TCR RLQ8 at concentrations of 0.78, 1.56, 3.12, 6.25, 12.5, 25, and 50.0 μM was injected over immobilized RLQ–HLA-A2 (1200 RU). (lower) Fitting curve for equilibrium binding that resulted in a $K_D$ of 9.7 μM. **(f)** (upper) TCR RLQ8 at concentrations of 0.78, 1.56, 3.12, 6.25, 12.5, 25, and 50.0 μM was injected over immobilized T1006I–HLA-A2 (1200 RU). (lower) Fitting curve for equilibrium binding that resulted in a $K_D$ of 122.4 μM.

**Figure S2.** **(a–b)** Electron density for the bound RLQ peptide in the two RLQ–HLA complexes in the asymmetric unit of the crystal. The $F_o - F_c$ omit map at 2.81 Å resolution is contoured at 1σ. The two RLQ peptides have identical conformations. **(c–d)** Electron density for the bound YLQ peptide in the two YLQ–HLA-A2 complexes in the asymmetric unit of the crystal. The $F_o - F_c$

omit map at 2.07 Å resolution is contoured at 1σ. The YLQ peptide displays two different conformations.

**Figure S3. (a)** Electron density at the interface in the RLQ3–RLQ–HLA-A2. Density from the final $2F_o - F_c$ map at 2.30 Å resolution is contoured at 1σ. **(b)** Electron density at the interface in the YLQ7–YLQ–HLA-A2 complex. Density from the final $2F_o - F_c$ map at 2.39 Å resolution is contoured at 1σ.

**Figure S4.** Comparison of Cα domain conformations in free versus ligand-bound TCR YLQ7. **(a)** The typical Cα conformation in free TCR YLQ7 (7N1D, this work). **(b)** The atypical Cα conformation in YLQ7 bound to YLQ–HLA-A2 (7N1F, this work). **(c)** The atypical Cα conformation in TCR 1F1E8hu (3MFF) (58). (**d**) CryoEM structure of the TCR–CD3 complex (6JXR) (59). The region of contact between Cα and the CD3δ subunit of the CD3εδ heterodimer is boxed. (**e**) Close-up of interactions between Cα and CD3δ. The side chains of contacting residues are drawn in stick representation.

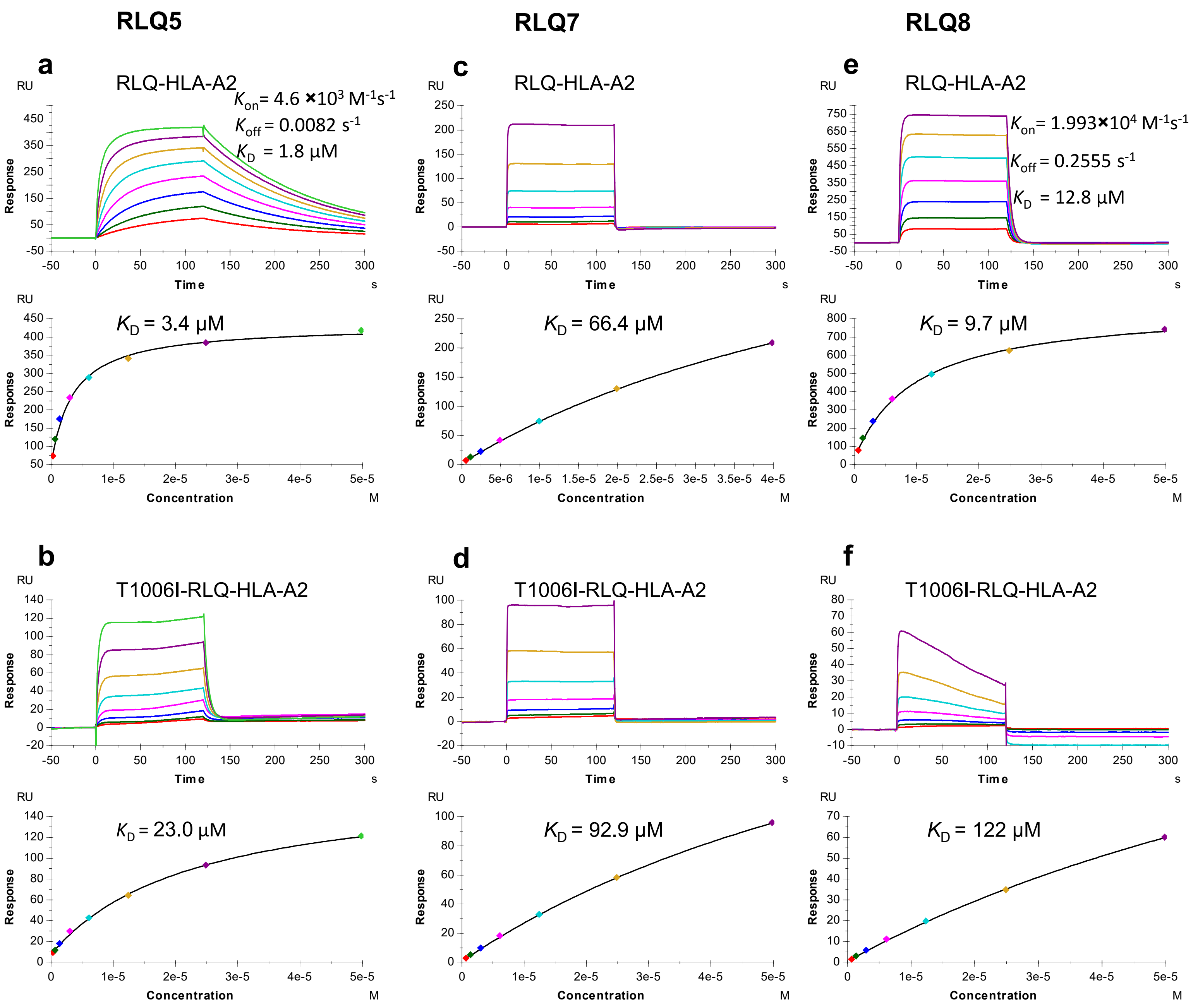
RLQ5
RLQ7
RLQ8
a
RLQ-HLA-A2
Kon= 4.6 ×10³ M⁻¹s⁻¹
Koff = 0.0082 s⁻¹
KD = 1.8 µM
KD = 3.4 µM
c
RLQ-HLA-A2
KD = 66.4 µM
e
RLQ-HLA-A2
Kon= 1.993×10⁴ M⁻¹s⁻¹
Koff = 0.2555 s⁻¹
KD = 12.8 µM
KD = 9.7 µM
b
T1006I-RLQ-HLA-A2
KD = 23.0 µM
d
T1006I-RLQ-HLA-A2
KD = 92.9 µM
f
T1006I-RLQ-HLA-A2
KD = 122 µM
RU
Response
Time
s
Concentration
M

Fig. S1

Fig. S2

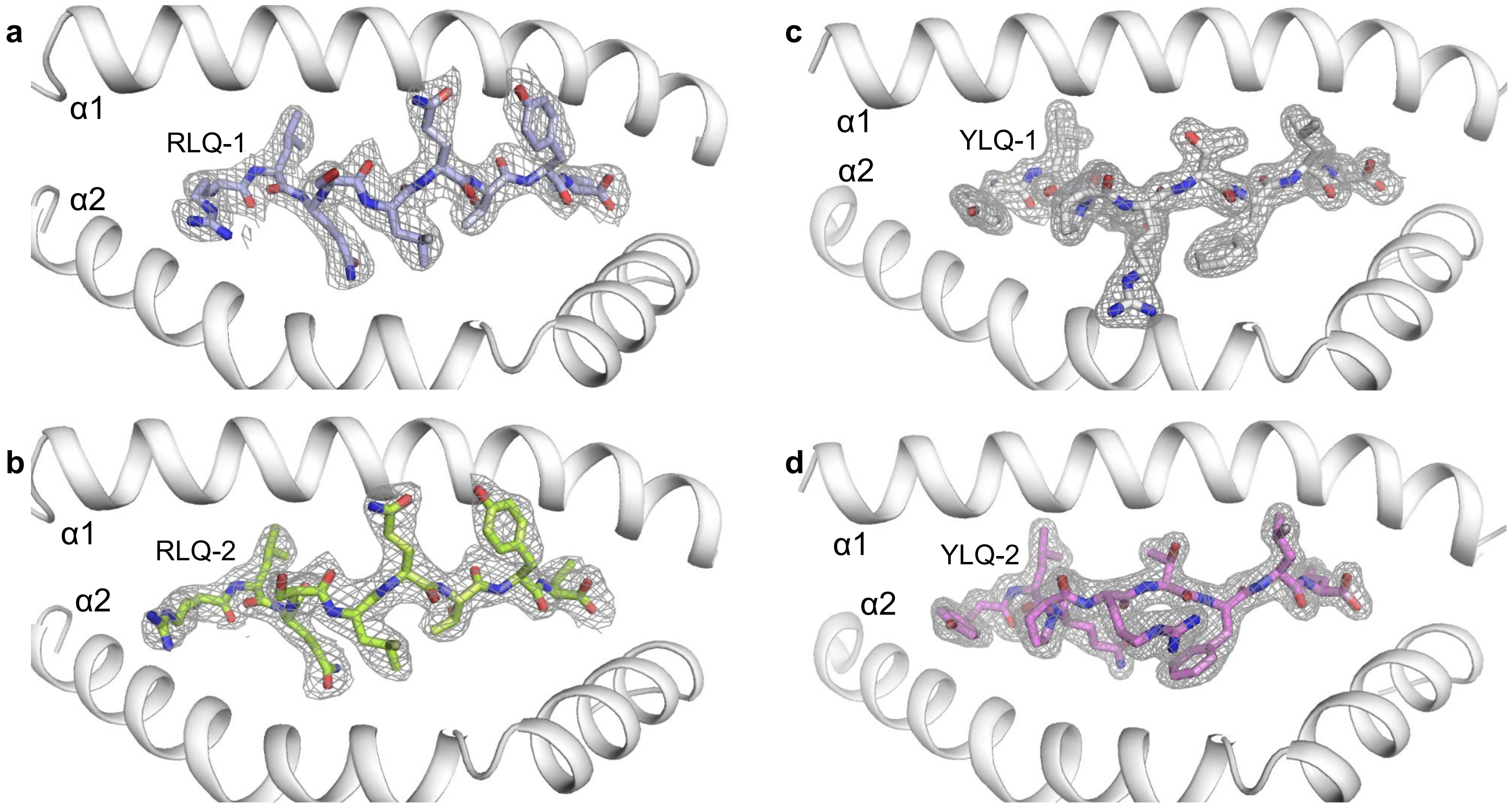

Fig. S3

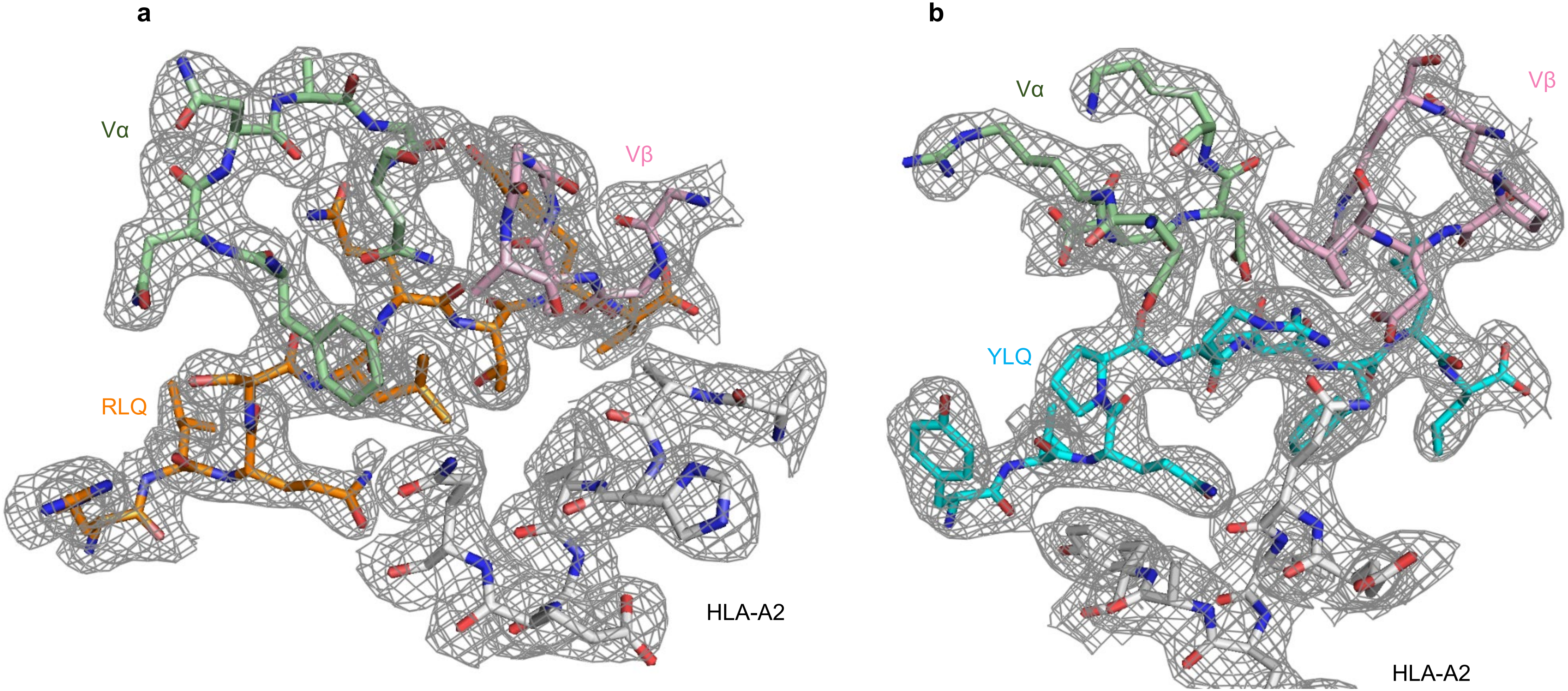

Fig. S4

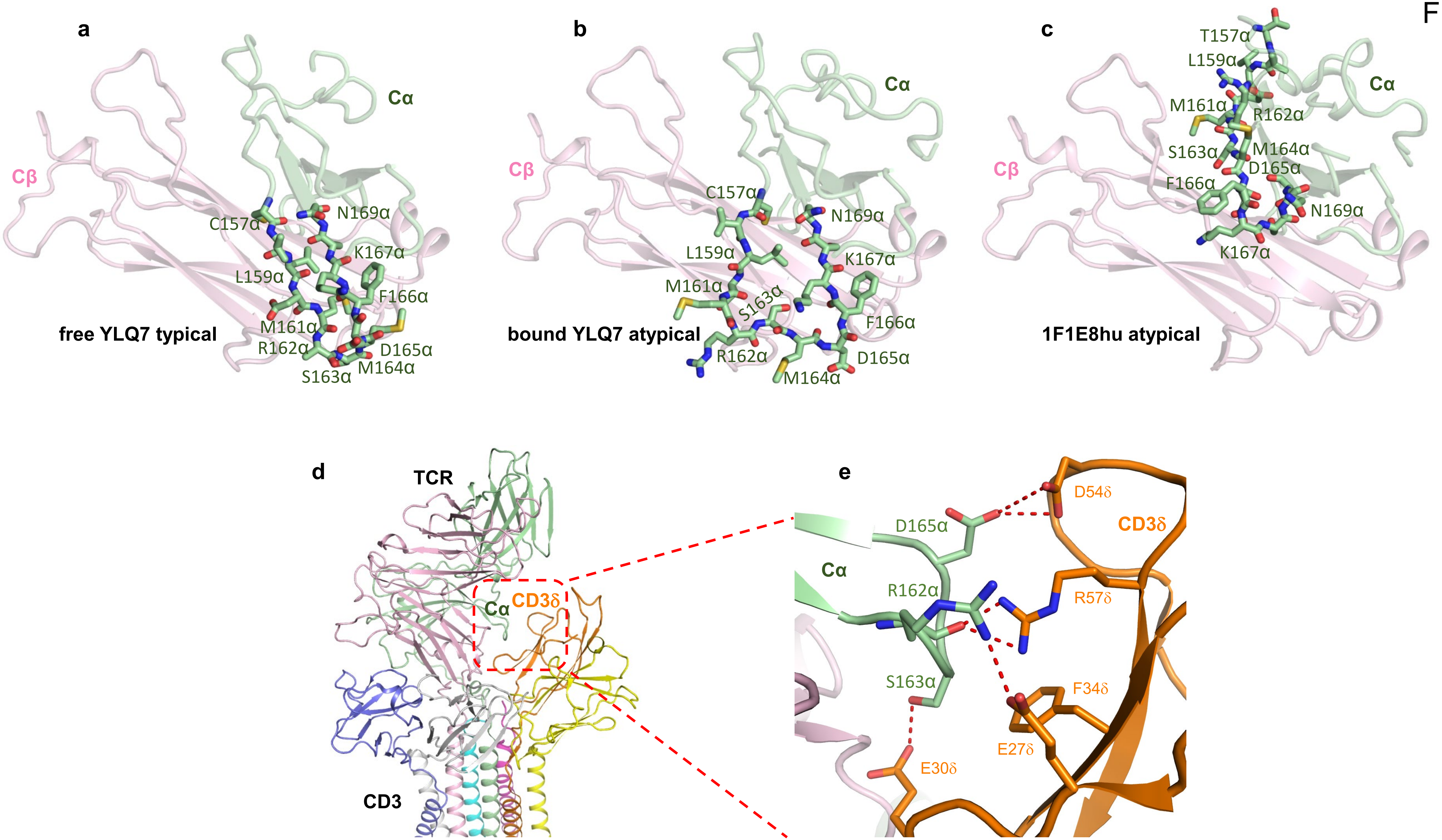

a
Cα
Cβ
C157α
N169α
K167α
L159α
F166α
free YLQ7 typical
M161α
R162α
D165α
S163α
M164α
b
Cα
Cβ
C157α
N169α
L159α
K167α
M161α
S163α
F166α
bound YLQ7 atypical
R162α
D165α
M164α
c
T157α
L159α
Cα
M161α
R162α
Cβ
S163α
M164α
D165α
F166α
N169α
K167α
1F1E8hu atypical
d
TCR
Cα
CD3δ
CD3
e
D54δ
D165α
CD3δ
Cα
R162α
R57δ
S163α
F34δ
E27δ
E30δ
